# Kingdom-wide evolutionary characterization of RNA editing factors across Archaeplastida

**DOI:** 10.1101/2025.10.09.681329

**Authors:** Ming Chen, Qingming Qu, Zhang Zhang

## Abstract

RNA editing, a post-transcriptional modification in plant mitochondria and plastids, is essential for environmental adaptation and diverse physiological processes. Despite extensive identification of RNA editing factors, primarily including pentatricopeptide repeat (PPR), multiple organelle RNA editing factor (MORF), organelle RNA recognition motif-containing (ORRM), and organelle zinc finger (OZ) proteins, their evolutionary history remains poorly understood. Here, we perform kingdom-wide evolutionary analyses across 364 high-quality Archaeplastida genomes and find massive PPR gene expansions in early-diverging land plants, predominantly driven by dispersed duplication associated with retroposition. Furthermore, integrative analyses indicate that DYW subgroup PPR genes have been horizontally transferred from plants to bdelloid rotifers, potentially enhancing the recipients’ adaptation to ultraviolet radiation. MORF proteins, accessory partners of PPRs, possess MORF hallmark domains structurally similar to protein-folding peptidase S8 propeptide/proteinase inhibitor I9 domains, suggesting a role in protein folding during RNA editing. Considering diverse domain compositions, we reclassify MORF, ORRM, and OZ proteins and uncover prevalent hallmark domain fusions. Together, these findings illuminate the kingdom-wide evolution of plant RNA editing machinery.

## Introduction

RNA editing, a post-transcriptional modification primarily involving cytidine-to-uridine (C-to-U) and uridine-to-cytidine (U-to-C) alterations in mitochondria and plastids, serves as a pivotal layer of genetic regulation in plants^1,2^. This process is essential for adaptation to diverse environments^3–6^ and various physiological processes^7^, including development^8^, growth^9^, flowering^10^, fruit ripening^11^, and resistance to stresses^12,13^. Editing factors, a group of nuclear-encoded proteins, are assembled into an editing complex to mediate RNA editing^6^. Given the remarkable diversity of editing factors in plants, elucidating deciphering their evolution is essential for understanding how the RNA editing machinery emerged and diversified^3,14–16^.

Decades of work have been made to identify RNA editing factors, primarily including Pentatricopeptide Repeat (PPR), Multiple Organelle RNA Editing Factor (MORF) (also known as RNA Editing Factor Interacting Protein, RIP), Organelle RNA Recognition Motif-containing (ORRM), and Organelle Zinc (OZ) finger proteins^14^, each characterized by hallmark domains or motifs that contribute distinctly to regulating RNA editing^17^. Among them, PPR proteins are key editing factors, specifically recognizing RNA sequences through P/PLS motif tracts and catalyzing C-to-U and U-to-C editing via carboxyl-terminal (C-terminal) DYW and DW:KP domains, respectively^18,19^. Furthermore, PPR proteins are classified into two subfamilies, P and PLS, with the latter further divided into PLS, E, E+, and DYW subgroups^17^. PPRs’ accessory proteins, including MORF, ORRM, and OZ, which are named after their MORF, RRM, Ran Binding Protein 2 (RanBP2) type Zinc finger (Znf) hallmark domains, respectively, coordinate RNA editing through interactions both with PPR proteins and among themselves^6,10,20–27^. Beyond their hallmark domains, these proteins also incorporate diverse additional domains, such as Glycine-Rich (GR) domain in MORF proteins^28–30^ and PPR/MORF domains in ORRM proteins^31^.

Building on these advances, studies have increasingly focused on the evolutionary histories of PPR, MORF, ORRM, and OZ genes^6,14,15,32,33^. The PPR gene family is one of the largest gene families in plants^19^, ranging from tens of members in early-diverging chlorophytes to hundreds or thousands in land plants^4,6^ and thereby reflecting massive expansions^4,34^. PPR genes also occur beyond plants, with P subfamily members widespread across eukaryotes and occasionally detected in prokaryotes^35,36^. Notably, DYW subgroup PPR genes are observed in multiple non-plant eukaryote lineages^37–40^, including Amoebozoa, Discoba, Fungi, Metazoa, and Malawimonadidae, presumably resulting from horizontal gene transfer (HGT)^41^. In contrast, MORF genes are present exclusively in angiosperms and gymnosperms^34^; ORRM genes are broadly distributed across Viridiplantae lineages including chlorophytes, mosses, liverworts, and angiosperms^31^; and OZ genes are sporadically distributed in gymnosperms, mosses, and lycophytes, but are absent from chlorophytes^25^. Despite these insights, a comprehensive evolutionary characterization of these RNA editing factors across Archaeplastida remains unexplored.

Toward this end, we collect a comprehensive list of genomes across Archaeplastida, systematically identify PPR, MORF, ORRM, and OZ genes, and reconstruct an evolutionary landscape of plant RNA editing factors. Building on the landscape, we perform phylogenomic analyses to elucidate the mechanisms underlying PPR expansions and investigate the extent and evolutionary significance of HGT in DYW subgroup PPR genes. Through sequence and structural alignments, we explore the conservation and diversity of hallmark domains of MORF proteins. Finally, we reclassify MORF, ORRM, and OZ proteins based on their domain composition.

## Results

### Evolutionary landscape of PPR, MORF, ORRM, and OZ genes across Archaeplastida

To systematically investigate the evolution of PPR, MORF, ORRM, and OZ genes, we obtain a total of 364 high-quality genomes across Archaeplastida that span early-diverging aquatic algae to highly derived land plants (for details see Materials and Methods), including glaucophytes (*n*=1), rhodophytes (*n*=7), prasinodermophytes (*n*=1), chlorophytes (*n*=33), charophytes (*n*=7), hornworts (*n*=22), mosses (*n*=13), liverworts (*n*=4), lycophytes (*n*=6), ferns (*n*=5), gymnosperms (*n*=13), and angiosperms (*n*=262) (Supplementary Table 1). Based on these genomes, we first identify 138 single-copy orthologous gene families (Supplementary Table 2) and then build a plant phylogenomic tree (Fig. 1a and Supplementary Fig. 1) that is topologically congruent with previous studies^42,43^. Next, we systematically identify 212,779 PPR, 2,759 MORF, 7,760 ORRM, and 829 OZ genes (Supplementary Tables 3 and 4), and evolutionarily characterize these genes in the phylogenomic tree as built above. Noticeably, we find that PPR genes are scarcely present in glaucophytes, rhodophytes, prasinodermophytes, chlorophytes, and charophytes, ranging only from 8 to 182 (Fig. 1b and Supplementary Table 4). In contrast, hundreds to thousands of PPR genes are detected in land plants, particularly some hornworts, mosses, and lycophytes, followed by gymnosperms, ferns, and angiosperms. Remarkably, two lycophytes, namely, *Selaginella moellendorffii* and *Isoetes sinensis*, have the highest number of PPR genes (*n*=3,861 and 3,256, respectively) among the 364 Archaeplastida species in our study (Fig. 1b and Supplementary Table 4). On the contrary, liverworts are short of PPR genes, less than 100 in count (Supplementary Table 4).

**Fig. 1:**
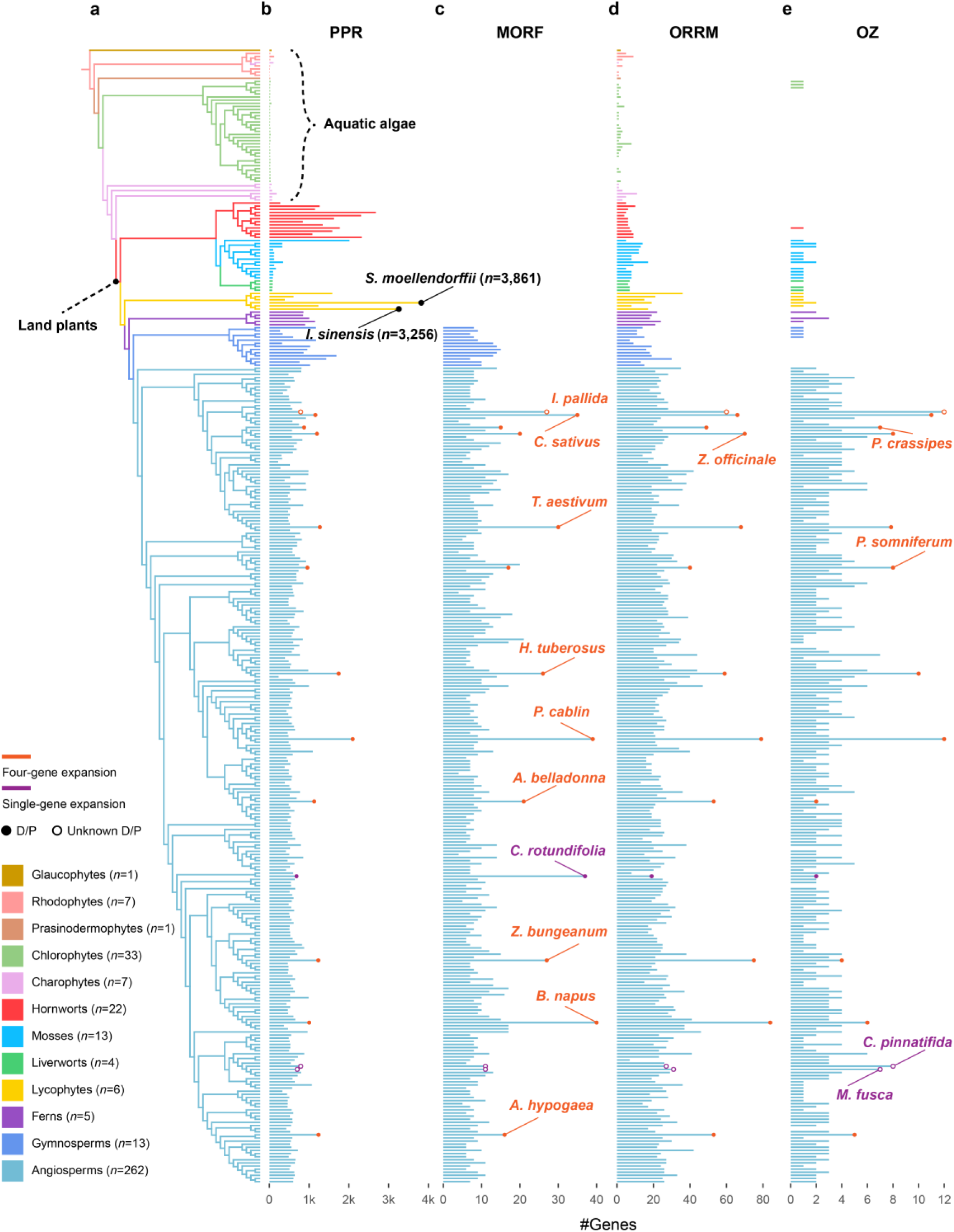
The evolutionary landscape of PPR, MORF, ORRM, and OZ genes across Archaeplastida. **(a)** The maximum likelihood phylogenomic tree of 364 Archaeplastida species, including glaucophytes (*n*=1), rhodophytes (*n*=7), prasinodermophytes (*n*=1), chlorophytes (*n*=33), charophytes (*n*=7), hornworts (*n*=22), mosses (*n*=13), liverworts (*n*=4), lycophytes (*n*=6), ferns (*n*=5), gymnosperms (*n*=13), and angiosperms (*n*=262). The best-fit model of each single-copy orthologous gene tree is automatically selected according to Bayesian Information Criterion in IQ-TREE. All gene trees (*n*=138) are merged and the final species tree is inferred by ASTRAL. Bootstraps are from 1,000 replicates. The distribution of **(b)** PPR, **(c)** MORF, **(d)** ORRM, and **(e)** OZ gene across Archaeplastida lineages. Two lycophytes, *S*. *moellendorffii* and *I*. *sinensis*, have the highest number of PPR genes (*n*=3,861 and 3,256, respectively). Orange and purple bars indicate gene expansions occurring simultaneously across all four gene families, and within individual families, respectively. Filled and open black circles denote species that have undergone genome diploidization/polyploidization (D/P), and those with unknown genome expansion, respectively. D/P: *Arachis hypogaea*, *Atropa bell adonna*, *B*. *napus*, *C*. *rotundifolia*, *C*. *sativus*, *Helianthus tuberosus*, *Papaver somniferum*, *P*. *cablin*, *Pontederia crassipes*, *T*. *aestivum*, *Zanthoxylum bungeanum*, and *Zingiber officinale*. Unknown D/P: *C*. *pinnatifida*, *Iris pallida*, and *M*. *fusca*.

Compared to PPR genes, the other three types of genes present distinct patterns. Specifically, MORF genes are found only in seed plants, including angiosperms and gymnosperms (Fig. 1c), in agreement with a previous study^34^. Additionally, seed plants generally harbor about 10 MORF genes, whereas some species, such as *Brassica napus*, *Pogostemon cablin*, and *Cissus rotundifolia*, have more MORF genes (*n*=40, 39, and 37, respectively; Supplementary Table 4). Conversely, ORRM genes are ubiquitous across Archaeplastida and exhibit a gradual increase from glaucophytes to angiosperms, indicating their ancient origin (Fig. 1d). Albeit much fewer in number than the rest three types of genes, OZ genes are enriched in angiosperms, with a minor presence in chlorophytes, hornworts, mosses, liverworts, lycophytes, ferns, and gymnosperms (Fig. 1e). Besides, many angiosperms that have undergone diploidization or polyploidization, such as *P. cablin*^44^, *Crocus sativus*^45^, and *Triticum aestivum*^46^, exhibit concurrent expansions of PPR, MORF, ORRM, and OZ genes (Fig. 1 and Supplementary Table 4). In contrast, among these four types of genes, MORF genes are the only ones markedly expanded in drought-resistant *C. rotundifolia* relative to other angiosperms, whereas OZ genes are the only ones significantly expanded in *Crataegus pinnatifida* and *Malus fusca*. Overall, PPR genes exhibit dramatic expansions in land plants, in contrast to MORF, ORRM, and OZ genes.

### PPR gene expansions in early-diverging land plants

Gene expansion can be achieved by gene duplication with four different modes, viz., whole genome duplication (WGD), tandem duplication (TD), proximal duplication (PD), and dispersed duplication (DD), which are classified according to genomic loci of duplicated gene pairs^47–49^. To explore the underlying mechanisms of PPR gene expansions in early-diverging land plants, we select high-quality chromosome-level genomes, including four hornworts (*Anthoceros agrestis*, 2,308 PPRs; *Notothylas orbicularis*, 1,772 PPRs; *Paraphymatoceros pearsonii*, 1,583 PPRs; and *Phymatoceros phymatodes*, 1,627 PPRs), one moss (*Takakia lepidozioides*, 2,016 PPRs), and two lycophytes (*I*. *sinensis*, 3,256 PPRs; and *Huperzia asiatica*, 1,583 PPRs) (Supplementary Table 4), with the charophyte *Zygnema circumcarinatum* (67 PPRs) serving as the phylogenetic outgroup. Furthermore, we select well-annotated PPR genes, accounting for 93.40∼100% of our identified PPRs, and detect that 90.04∼95.89% of them form duplicated gene pairs based on sequence similarity analysis (Supplementary Table 5). Meanwhile, we infer WGDs in these selected plants through comprehensive analyses of gene synteny and synonymous substitution rates (Ks) of homologous gene pairs (for details see Materials and Methods). Accordingly, we find that *Z. circumcarinatum* and four hornworts lack evidence of WGD, whereas *T. lepidozioides* and *I. sinensis* each experience one WGD, and *H. asiatica* undergoes two WGDs (Supplementary Fig. 2-9). These results are consistent with previous studies^5,50–54^, except that *I. sinensis* has been reported to experience two WGDs based solely on Ks distributions^55^. Consequently, integration of inferred WGD events with comparison of genomic loci of duplicated PPR gene pairs (for details see Materials and Methods) indicates that DD is primarily responsible for PPR gene expansions in the selected species, followed by WGD in *I. sinensis* and *H. asiatica* (Fig. 2a and Supplementary Table 5).

**Fig. 2:**
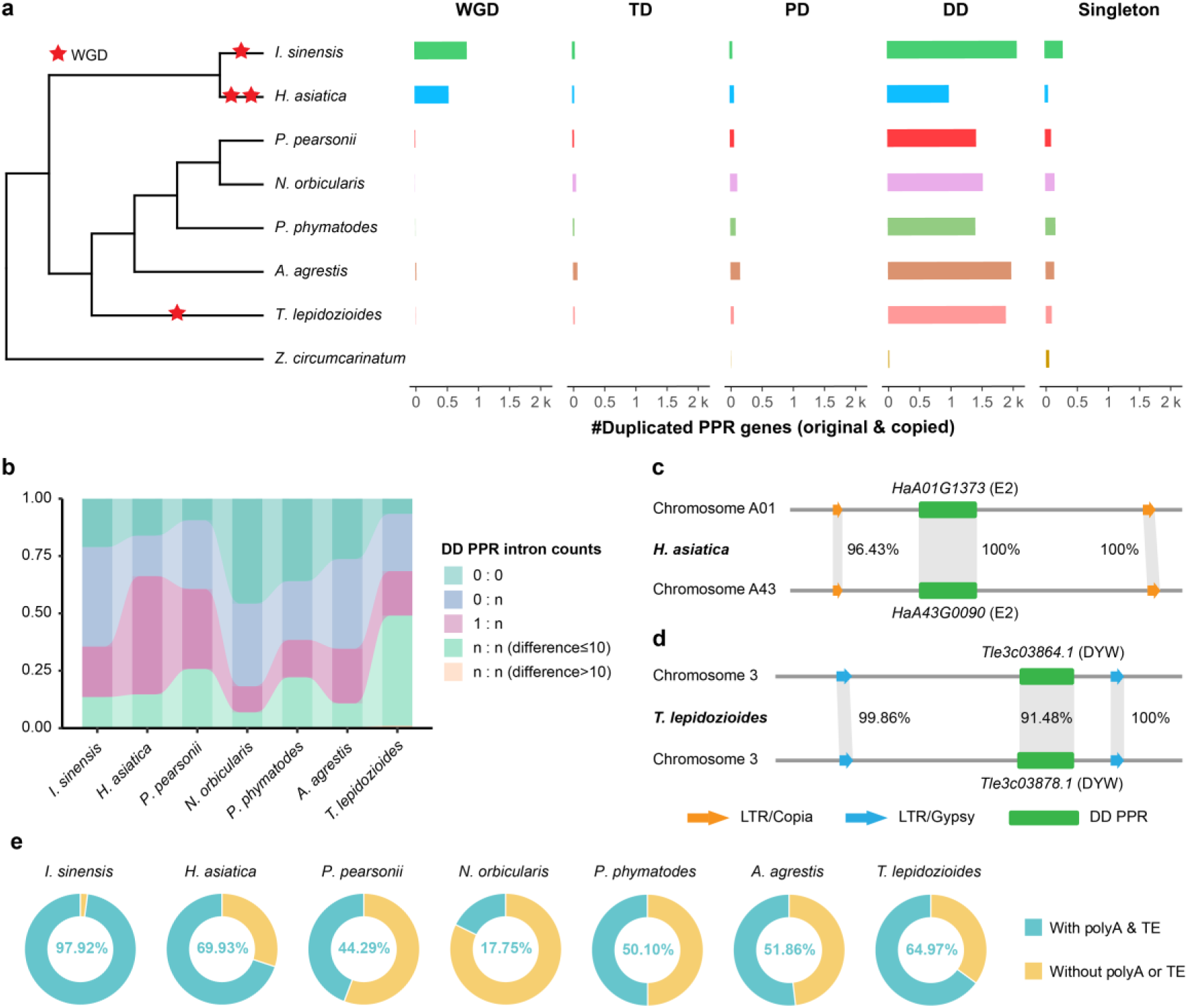
Mechanisms of PPR gene expansions. **(a)** Numbers of duplicated PPR genes (original and copied) derived from WGD, TD, PD, and DD in seven representative species (*A*. *agrestis*, *N*. *orbicularis*, *P*. *pearsonii*, *P*. *phymatodes*, *T*. *lepidozioides*, *I*. *sinensis*, and *H*. *asiatica*) and the outgroup (*Z*. *circumcarinatum*). A red five-pointed star denotes a whole genome duplication event. WGD: Whole Genome Duplication, TD: Tandem Duplication, PD: Proximal Duplication, DD: Dispersed Duplication. **(b)** Percentage of intronless DD PPR gene pairs. Numbers to the left and right of the colon indicate intron counts in each DD PPR gene pair. Categories “0:0”, “0:n”, “1:n”, and “n:n (difference > 10)” denote intron loss in DD PPR gene pairs. **(c)** E2 and **(d)** DYW subgroup PPR gene pairs (DD) flanked by similar LTR/Copia and LTR/Gypsy, respectively. **(e)** Percentages of intronless DD PPR gene pairs with polyA tails and flanked by LTRs.

Furthermore, it has been reported that DD is related to retroposition^47,49^, a process that the complementary DNA transcript of a gene is inserted back into the genome, thus producing a retrogene^56^. Generally, the retrogene of a duplicated pair features intron loss and a polyadenylation (polyA) tail, and the pair is flanked by short direct repeats (e.g., transposable element, TE)^56,57^. Thus, we examine these features of DD PPR gene pairs as indicators of retroposition. Specifically, we compare intron counts of PPR genes in each dispersed duplicate pair, and find that 74.78∼93.21% of DD-derived PPR genes in hornworts and lycophytes have undergone intron loss, followed by the moss *T. lepidozioides* with 51.98% (Supplementary Table 6 and Fig. 2b; for details see Materials and Methods). Furthermore, we investigate the presence of polyA tails downstream of the 3’ untranslated regions (3’UTRs) of intronless DD-derived PPR genes and find that 97.99∼99.26% possess polyA tails in lycophytes, 70.37∼75.61% in hornworts, and 81.55% in the moss *T. lepidozioides* (Supplementary Fig. 10). Meanwhile, we identify TEs within 20 kilobases (kb) upstream and downstream of intronless DD PPR gene pairs, showing that 70.52∼99.93% of them are flanked by TEs in lycophytes, 26.56∼68.31% in hornworts, and 80.11% in *T. lepidozioides* (Supplementary Fig. 11). Notably, most of these TEs are long terminal repeats (LTRs), including LTR/Copia, LTR/Gypsy, and LTR/unknown (Supplementary Table 7). Additionally, we present two DD PPR gene pairs with intron loss that are flanked by the same type of LTR with high sequence similarity, viz., E2 subgroup PPR genes, *HaA01G1373* and *HaA43G0090* in *H*. *asiatica* flanked by LTR/Copia (96.43% upstream and 100% downstream) (Fig. 2c), and DYW subgroup PPR genes, *Tle3c03864.1* and *Tle3c03878.1* in *T. lepidozioides* by LTR/Gypsy (99.86% upstream and 100% downstream) (Fig. 2d). Furthermore, co-occurrence of these three retroposition features is detected in 69.93∼97.92% of DD PPR gene pairs in lycophytes, 17.75∼51.86% in hornworts, and 64.97% in *T. lepidozioides* (Fig. 2e), suggesting lineage-specific degeneration of retroposition signals. Collectively, these results indicate that PPR gene expansions in early-diverging land plants are primarily driven by DD associated with retroposition.

### Horizontal transfer of DYW subgroup PPR genes from plants to bdelloid rotifers

As PPR genes consist of the P and PLS subfamilies, we explore their phylogenetic patterns by reconstructing an evolutionary landscape across Archaeplastida (Supplementary Fig. 12). We find that the P subfamily PPR genes are ubiquitously distributed in Archaeplastida, with their numbers progressively increasing from early-diverging glaucophytes to highly derived angiosperms. Conversely, the PLS subfamily, including PLS, E, E+, DYW, and DYW:KP subgroups, is confined to land plants, with rare occurrence in liverworts and some mosses. In addition, the PLS and E subgroups are numerous in lycophytes, while the E+ subgroup is exclusively enriched in hornworts. The DYW subgroup is broadly distributed in land plants, with a maximum of 750 members observed in the moss *T. lepidozioides*. In contrast, its variant, the DYW:KP subgroup, is restricted to hornworts, lycophytes, and ferns (Supplementary Table 3 and Supplementary Fig. 12), conforming with a previous study^34^.

Inspired by the previous findings that DYW subgroup PPR genes have also been detected in non-plants^37,41^, we next investigate the evolutionary relationships of the DYW and DYW:KP subgroups between plants and non-plants. Given that repetitive and variable PLS motif triplets may hinder reliable phylogenetic inference and DYW domain catalyzes C-to-U editing via its PG box (covering the first two β-strands), the zinc ion binding signature HxE(x)_n_CxxC, and C-terminal DYW tripeptide^58–60^, we extract 6,646 DYW and 3,761 DYW:KP domains of 68 land plants as representatives of their respective subgroup PPR genes (Supplementary Table 8). Based on these plant domains, we further survey non-plant DYW and DYW:KP domains across 40 lineages spanning eukaryotes, prokaryotes, and viruses (Supplementary Table 9; for details see Materials and Methods). In total, we detect 1,666 DYW and 6 DYW:KP domains in Metazoa, 31 DYW in Rhizaria, 22 DYW and 1 DYW:KP in Discoba, 15 DYW in Fungi, 9 DYW in Alveolata, 3 DYW in Haptista, and 1 DYW in Amoebozoa (Supplementary Table 10), exhibiting 21.21∼80% sequence similarity to their plant counterparts (Supplementary Table 11). To reduce domain redundancy, we further select 1,151 DYW and 356 DYW:KP domains from six representative species spanning major land plant lineages, including *P. pearsonii* (hornwort), *T. lepidozioides* (moss), *I. sinensis* (lycophyte), *Alsophila spinulosa* (fern), *Gnetum montanum* (gymnosperm), and *Amborella trichopoda* (angiosperm), together with all non-plant DYW and DYW:KP domains for phylogenetic reconstruction (Supplementary Table 10). Notably, three major clusters of non-plant DYW domains are embedded within plant lineages, suggesting multiple waves of HGT from plants to non-plants (Fig. 3).

**Fig. 3:**
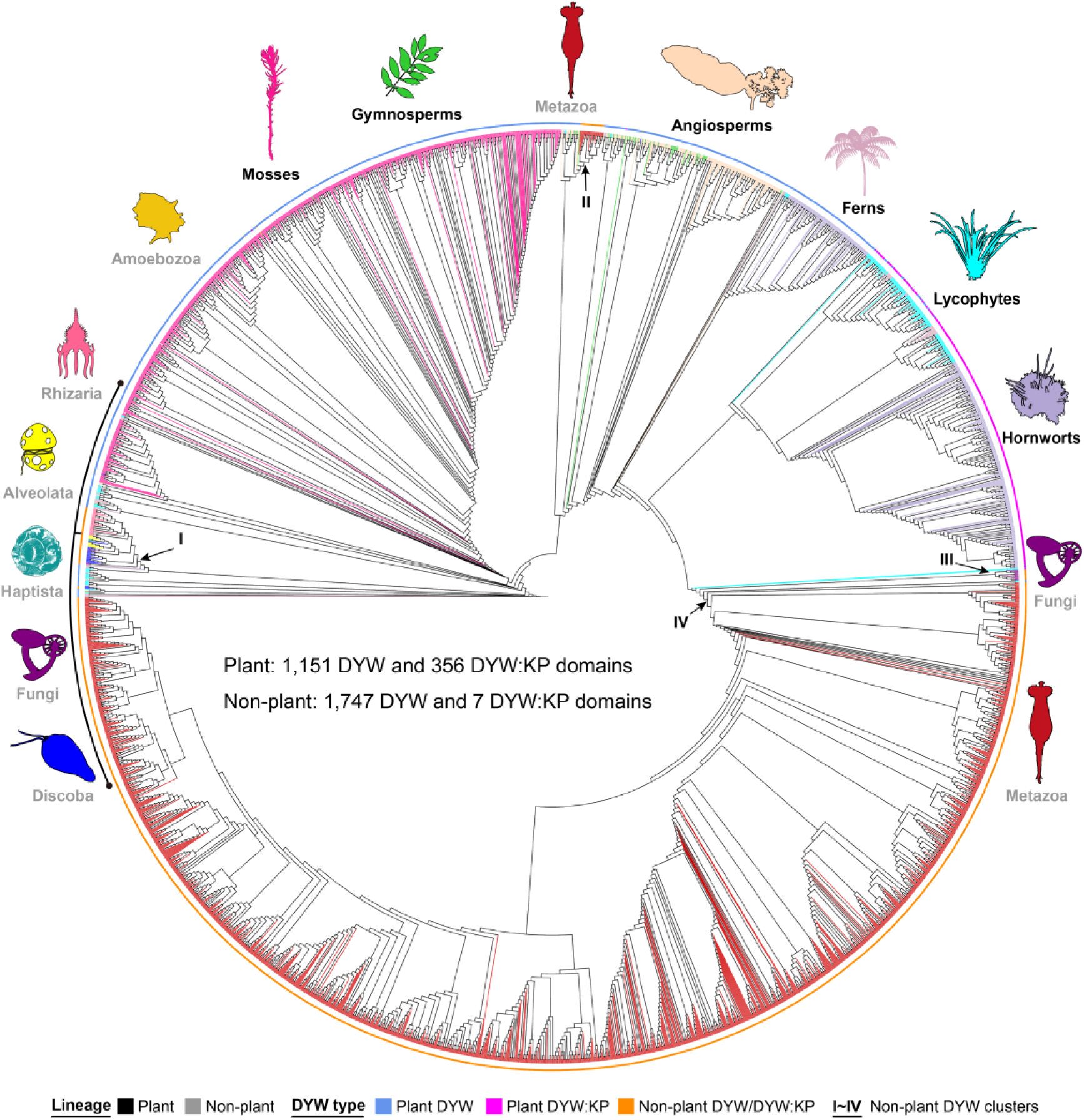
The maximum likelihood phylogenetic tree of DYW and DYW:KP domains in plants and non-plants. Three predominant clusters (I, II, and III) of non-plant DYW domains are embedded within plants. Among non-plant lineages, Metazoa contain the largest number of DYW domains. Best-fit model (IQ-TREE): JTT + R10. Bootstraps are from 1,000 replicates.

To explore the potential role of DYW subgroup PPR genes in non-plants, we select a representative DYW domain from each lineage and compare its sequence and structure with those of plant donors. On the one hand, most of these non-plant DYW domains contain the conserved HxE(x)nCxxC motif, except for the HxE(x)nGxxN in Fungi, whereas the PG-box (approximately 15 amino acid residues (aa) beginning with proline and glycine^61^) and DYW tripeptide are variable (Fig. 4a and 4b; Supplementary Fig. 13). On the other hand, these non-plant DYW domains exhibit significant structural similarity to those of their plant donors (Root Mean Square Deviation, RMSD ≤ 2.5Å^62^), such as the DYW domain in the bdelloid rotifer *Adineta vaga* (Metazoa) and its counterpart in the moss *T. lepidozioides* (Fig. 4b and Supplementary Fig. 13). Surprisingly, Metazoa acquire the largest number of DYW domains from plants (Fig. 3), all of which occur in bdelloid rotifers that are resistant to ultraviolet (UV) radiation and inhabit diverse environments, such as mosses, lichens, soils, and deserts^63,64^. Notably, these bdelloid rotifers comprise *A. vaga*, *A*. *ricciae*, *A. steineri*, *Didymodactylos carnosus*, *Philodina roseola*, *Rotaria magnacalcarata*, *R. socialis*, *R. sordida*, *R.* sp. ‘Silwood1’, and *R.* sp. ‘Silwood2’. Among them, we find that *A. vaga* harbors lots of PPR genes (*n*=191), nearly one third of which (*n*=60) are members of the DYW subgroup (Supplementary Table 12). The majority of these DYW subgroup PPR genes (58/60) are expressed (Fig. 4c), suggesting their potential roles in biological processes. Intriguingly, the moss *T. lepidozioides*, a potential donor of DYW subgroup PPR genes to *A. vaga*, is also resistant to UV radiation^5^. Consistent with this resistance, its DYW genes are upregulated during recovery from UV irradiation (Fig. 4d and Supplementary Table 13), highlighting their potential roles in response to UV radiation. Taken together, these results suggest that bdelloid rotifers living in mosses may have acquired the DYW subgroup PPR genes as an adaptation to high UV radiation (Fig. 4e).

**Fig. 4:**
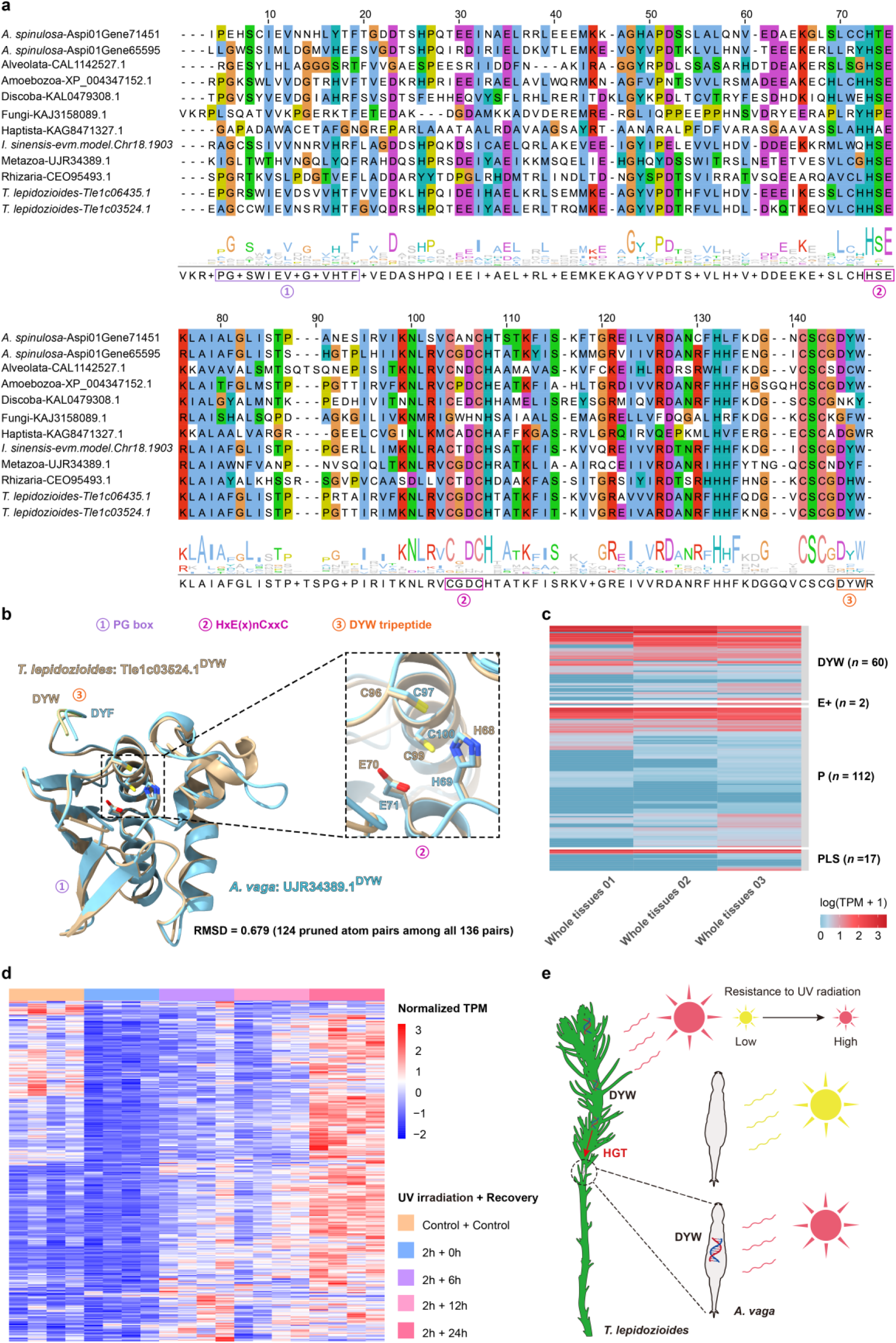
HGT of DYW domains from plants to bdelloid rotifers. (a) Sequence alignments between plant and non-plant DYW domains. Amino acid residues in rectangles denote PG box, HxE(x)nCxxC, and DYW tripeptide. **(b)** Protein structure alignment between DYW domain (protein ID: Tle1c03524.1) in the moss *T. lepidozioides* and its counterpart (protein ID: UJR34389.1) in the bdelloid rotifer *A. vage*. **(c)** Expression of PPR genes (DYW, E+, PLS, and P subgroups) in whole tissues of *A. vage*. TPM: Transcripts Per Million. **(d)** Expression of DYW subgroup PPR genes in *T. lepidozioides* under control condition and after 2-hour UV irradiation, followed by 0, 6, 12, and 24-hour recovery. **(e)** A proposed model of horizontal transfer of DYW subgroup PPR genes from mosses (e.g., *T. lepidozioides*) to bdelloid rotifers (e.g., *A. vage*).

### Evolutionary conservation and diversity of MORF domains

MORF proteins, exclusive to seed plants (Fig. 1c) and comprising nine members (MORF1∼MORF9) in *A. thaliana*, are characterized by their MORF hallmark domains and are essential for PPR-mediated RNA editing^28–30^. To explore the evolutionary conservation of MORF domains, we extract 334 instances from 33 representative seed plants (Supplementary Table 14) and identify their homologs across 45 lineages spanning eukaryotes, prokaryotes, and viruses (Supplementary Table 9). We find that MORF domains are homologous to peptidase S8 propeptide/proteinase inhibitor I9 domains (also known as PA domain^28^) in metazoa, fungi, bryophytes, lycophytes, and ferns (Fig. 5a and Supplementary Table 15). Furthermore, we compare the structures of the MORF domain and its homologous PA domain with the highest sequence similarity in each of these five lineages. We find that although they have relatively low sequence similarity (33.33∼41.94%), their structures are significantly similar (RMSD ≤ 2.5Å), with RMSD values varying from 0.705∼1.039Å (Fig. 5b and 5c; Supplementary Fig. 14). Notably, MORF domains from MORF3 and MORF8 proteins share the same α-helix and β-strand architecture as PA domains (Fig. 5c and Supplementary Fig. 14). As PA domain is important in assisting fold zymogens and mature peptidases^65,66^, these results imply that MORF domains from MORF3 and MORF8 proteins may help fold PPR proteins to regulate RNA editing.

**Fig. 5:**
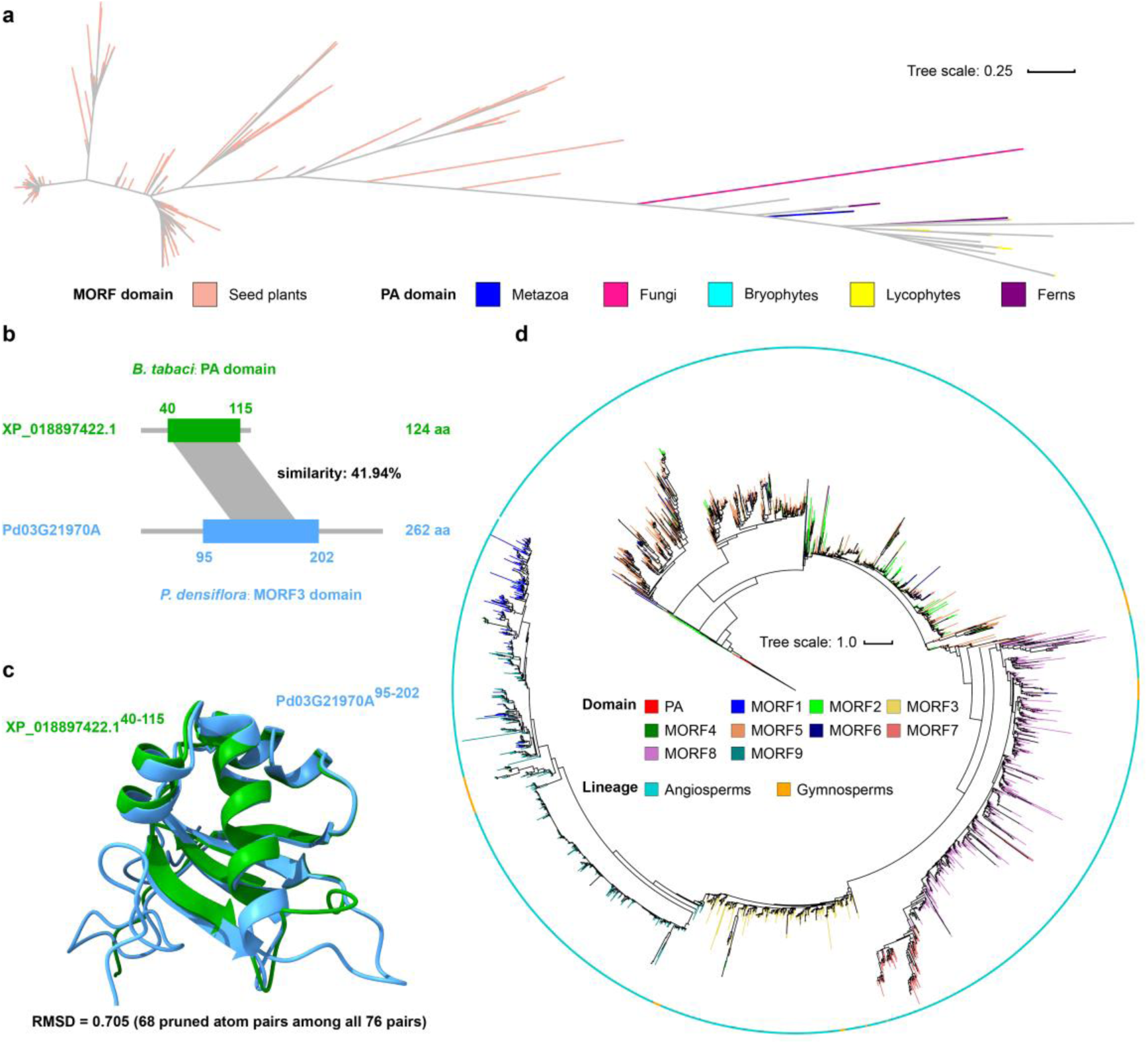
Conservation and diversity of MORF domains. **(a)** The maximum-likelihood phylogenetic tree of MORF and PA domains. The tree scale bar represents the number of substitutions per site. Model (FastTree): Jones-Taylor-Thorton, CAT approximation with 20 rate categories. **(b)** Sequence and **(c)** structure alignments between MORF3 domain (protein ID: Pd03G21970A) from *Pinus densiflora* and its homologous PA domain (protein ID: XP_018897422.1) from *Bemisia tabaci*. **(d)** The maximum-likelihood phylogenetic tree of MORF1∼MORF9 domains from angiosperms and gymnosperms. Five non-seed plant PA domains with highest sequence similarity to MORF domains are used as the outgroup (protein IDs: Dicom.22G070000.1.p, HaB36G042, Phcar.S3G189600.t1, Phsp.C2G192800.t1, and XP_024533311.1). Best-fit model (IQ-TREE): JTT + R8. Bootstraps are from 1,000 replicates.

To investigate the evolutionary diversity of MORF domains, we classify them into MORF1 to MORF9 as identified in *A. thaliana* and survey PA domains across our collected 364 high-quality Archaeplastida genomes. Accordingly, PA domains are ubiquitous across major lineages, including glaucophytes, charophytes, mosses, liverworts, hornworts, lycophytes, ferns, gymnosperms, and angiosperms (Supplementary Fig. 15). Furthermore, we reconstruct a phylogenetic tree of the 334 MORF domains, using five non-seed plant PA domains with the highest sequence similarity as the outgroup. Consequently, MORF2, MORF5, and MORF6 domains cluster into early-diverging clades in the phylogenetic tree, indicating their earlier origins relative to other MORF domains in seed plants (Fig. 5d). Moreover, most MORF domains (MORF3∼6, 8, and 9) are broadly distributed across angiosperms and gymnosperms, suggesting that MORF diversification predates the major divergences among seed plant lineages. At the gene level, we analyze the expression of MORF1∼MORF9 genes, each harboring a single MORF hallmark domain, across multiple tissues of the gymnosperm *Torreya grandis* and the angiosperm *A. thaliana*. Notably, among these MORF genes, MORF8 and MORF9 display higher expression across tissues in both species, consistent with observations in *Apium graveolens* and *Populus trichocarpa*^67,68^, indicating their conservation in expression (Supplementary Fig. 16).

### Hallmark domain fusions among MORF, ORRM, OZ, and PPR proteins

While MORF proteins contain the MORF hallmark domain, the presence of additional domains, such as the GR domain in MORF1, MORF4, and MORF8 proteins^29^, suggests the novel domain composition and underscores the need for an updated classification of MORF proteins. Thus, in terms of domain composition, we classify our identified 2,759 MORF proteins into four groups:

1. Single-Pure MORF (SP-MORF, *n*=1,916), containing a single MORF domain;
2. Single-Mixed MORF (SM-MORF, *n*=544), containing a single MORF domain plus one or more non-MORF domains; (3) Multiple-Pure MORF (MP-MORF, *n*=42), containing multiple MORF domains; and (4) Multiple-Mixed MORF (MM-MORF, *n*=257), containing multiple MORF domains plus one or more non-MORF domains (Fig. 6a and Supplementary Table 16).

**Fig. 6:**
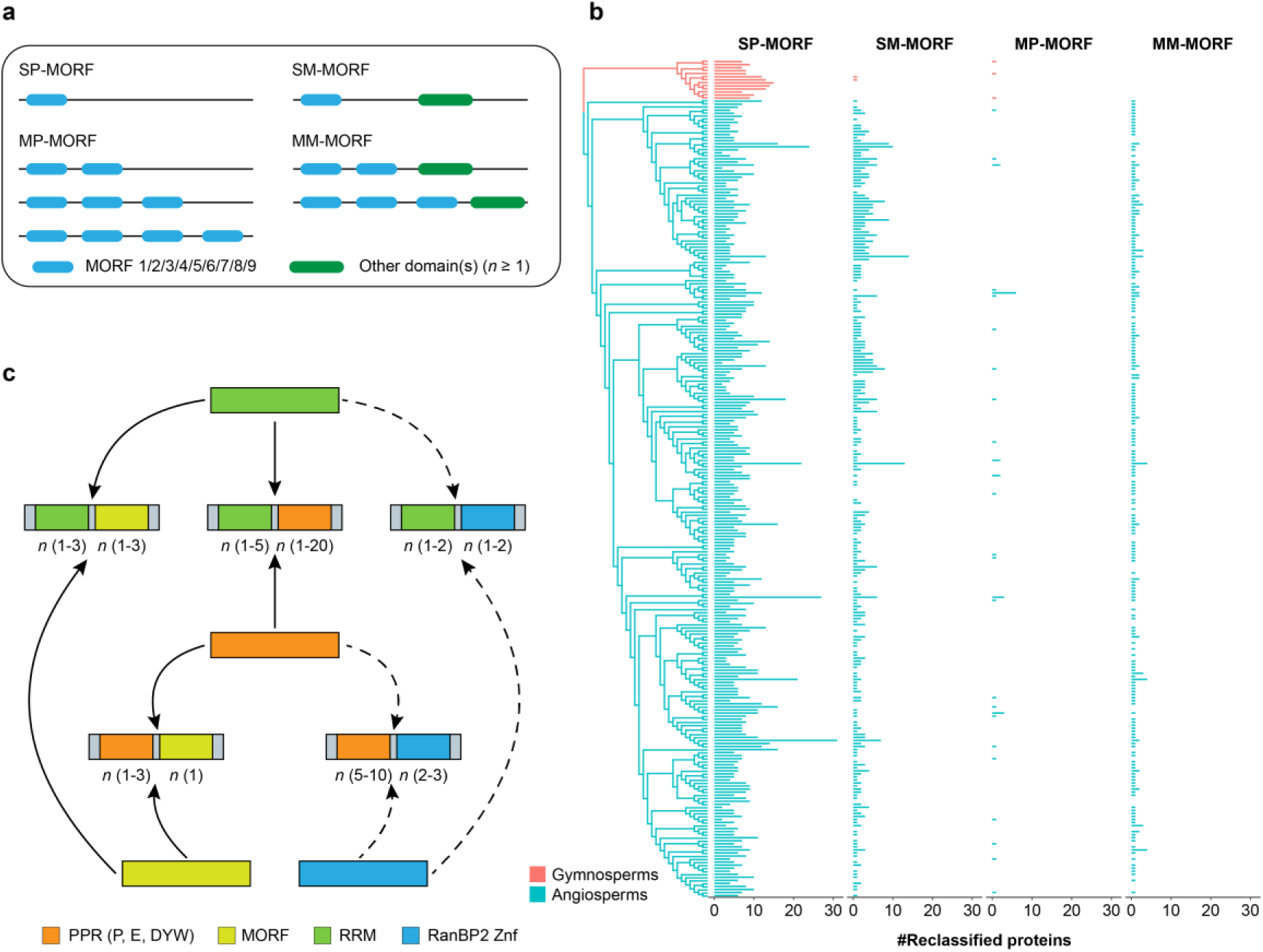
Reclassification and diversity of PPR accessory proteins. **(a)** Reclassification of MORF proteins based on domain composition. Blue and green bars represent MORF1∼MORF9 and other domains, respectively. SP-MORF: Single-Pure MORF, SM-MORF: Single-Mixed MORF; MP-MORF: Multiple-Pure MORF; MM-MORF: Multiple-Mixed MORF. **(b)** The phylogenetic tree of reclassified MORF proteins from angiosperms and gymnosperms. **(c)** Fusions among PPR, MORF, RRM, and RanBP2 Znf domains. Numbers in parentheses indicate domain counts. Proteins marked with solid and dashed arrows are localized to mitochondria/plastids and other subcellular compartments, respectively.

Furthermore, we reconstruct the evolutionary landscape of these reclassified MORF proteins across 262 angiosperms and 13 gymnosperms (Fig. 6b). Specifically, SP-MORF proteins are widespread in gymnosperms and angiosperms with larger numbers, whereas SM-MORF proteins are abundant in angiosperms but rare in gymnosperms. Conversely, MP-MORF proteins occur sparsely in both lineages and MM-MORF proteins are only observed in angiosperms. Importantly, these reclassified MORF proteins exhibit a wide range of domain compositions. Among them, 492 SM-MORF and 9 MM-MORF proteins harbor a GR domain alongside MORF1∼6, 8, and 9 domains in angiosperms, whereas in the gymnosperm *T*. *grandis*, a single SM-MORF protein contains a GR domain co-occurring with MORF9 domain, extending the previously observed MORF-GR domain architectures in MORF1, MORF4, and MORF8 proteins^29^. Surprisingly, we find that MORF domains and P motifs are fused in two SM-MORF proteins, viz., AY_017782_RA in the angiosperm *Aquilaria yunnanensis* and KAH9320946.1 in the gymnosperm *Taxus chinensis* (Supplementary Table 16). Moreover, MORF domain and the RRM hallmark domain of ORRM are fused in SM- and MM-MORF proteins in angiosperms. Specifically, these domain fusions fall into four types: 1-MORF & 1-RRM (*n*=23), 1-MORF & 3-RRM (*n*=1), 2-MORF & 1-RRM (*n*=243), 3-MORF & 1-RRM (*n*=3) (Supplementary Fig. 17 and Supplementary Table 16), where the leading number indicates the copy number of each corresponding domain. Notably, the most abundant fused type, 2-MORF & 1-RRM, is represented by the RNA editing factor ORRM1^69^.

ORRM proteins, specifically, ORRM1 to ORRM6, have been identified in several angiosperms as mediators of RNA editing^23,70^. Likewise, we reclassify our identified 7,760 ORRM proteins into SP-ORRM (*n*=2,689), SM-ORRM (*n*=1,751), MP-ORRM (*n*=2,948), and MM-ORRM (*n*=372) (Supplementary Table 17). Furthermore, we reconstruct their evolutionary landscape across Archaeplastida, and find that SP-, SM-, and MP-ORRM proteins exhibit a gradual increase from glaucophytes to angiosperms with larger number, while MM-ORRM proteins show a sparse distribution (Supplementary Fig. 18). In addition to previously identified ORRM1 to ORRM6 with a single RRM domain, MP-ORRM proteins, such as CP29A, CP31A, and CP31B, each containing two RRM domains, also affect RNA editing^23^. Intriguingly, numerous RRM and PPR domains are fused in SM- and MM-ORRM proteins across Archaeplastida (Supplementary Fig. 19). Specifically, SM-ORRM proteins contain the highest number of RRM-P fusions in angiosperms (*n*=171), followed by hornworts (*n*=6), mosses (*n*=4), liverworts (*n*=4), charophytes (*n*=3), gymnosperms (*n*=3), lycophytes (*n*=1), and ferns (*n*=1). Moreover, SM-ORRM proteins exhibit one RRM-DYW fusion in hornworts and angiosperms, and three RRM-E fusions in angiosperms (Supplementary Table 17). Conversely, MM-ORRM proteins have only five RRM-P and one RRM-E fusions in angiosperms. In addition, fusions of RRM and the RanBP2 Znf hallmark domain of OZ protein are detected in proteins that are not targeted to mitochondria or plastids (Supplementary Table 18).

Similarly, we reclassify OZ proteins and explore their evolution using the same approach applied to MORF and ORRM proteins. Consequently, we identify 761 MP-OZ, 61 MM-OZ, 6 SP-OZ, and 1 SM-OZ proteins (Supplementary Table 19). Based on their evolutionary landscape across Archaeplastida, we find that MP-OZ proteins are enriched in angiosperms (*n*=725), followed by mosses (*n*=14), lycophytes (*n*=6), ferns (*n*=6), gymnosperms (*n*=4), chlorophytes (*n*=3) and liverworts (*n*=3) (Supplementary Fig. 20). Additionally, 60 MM-OZ, 6 SP-OZ, and 1 SM-OZ proteins are found in angiosperms, while a single MM-OZ occurs in hornworts. Notably, we detect that RanBP2 Znf and PPR domains (P and E) are fused in four angiosperm proteins, despite their absence of targeting to mitochondria or plastids (Supplementary Table 20). Collectively, our results indicate that hallmark domains of PPR, MORF, ORRM, and OZ proteins can be fused with more diverse patterns than previously thought (Fig. 6c).

## Discussion

Our kingdom-wide characterization of RNA editing factors across Archaeplastida reveals a complex evolutionary trajectory defined by massive gene expansions, HGT, and the diversity and fusion of hallmark domains.

PPR genes, the primary mediators of plant RNA editing, exhibit dramatic expansions in hornworts, mosses, and lycophytes. We demonstrate that these expansions are predominantly driven by dispersed duplication associated with retroposition, evidenced by signatures such as intron loss, polyA tails, and flanking TEs. In contrast, WGD plays a subsidiary role in PPR expansion in lycophytes and contributes little to that in the moss *T. lepidozioides* (Fig. 2a). Notably, *T. lepidozioides* exhibits predominant dispersed duplication but limited retroposition features, suggesting large-scale gene fractionation after an ancient WGD event^71^. These results imply that retroposition, as well as WGD, provided the genomic plasticity necessary for early land plants to diversify their RNA editing machinery during the colonization of harsh terrestrial environments^4^.

The evolutionary reach of plant RNA editing extends far beyond Archaeplastida. The discovery of DYW subgroup PPR genes in diverse non-plant lineages—including Metazoa, Amoebozoa, Fungi, Haptista, and Rhizaria—indicates that DYW-mediated C-to-U editing is a broadly distributed eukaryotic mechanism, which is more common than previously recognized^41,72,73^. Noticeably, the high expression of these genes in both recipient bdelloid rotifers (Metazoa) and their putative plant donors (e.g., *T. lepidozioides*), coupled with shared resistance to UV radiation, suggests a convergent adaptation where RNA editing may facilitate the repair of UV-induced damage^74,75^. This highlights the functional redeployment of editing factors following cross-kingdom transfer to meet specific environmental pressures.

MORF proteins, restricted to seed plants, display significant structural similarity between their hallmark MORF domains and PA domains of other proteins. Specifically, MORF3 and MORF8 domains more closely resemble PA domains associated with protein folding, whereas MORF1 domain has been reported to be more structurally similar to an N-terminal ferredoxin-like domain that confers RNA substrate positioning in bacterial 4-thio-uracil tRNA synthetases^28^. Our results suggest that the structural diversity of MORF domains underpins their functional versatility in RNA editing regulation.

Prompted by the structural diversity observed among MORF domains, we extended our analysis to MORF, ORRM, and OZ proteins, systematically reclassifying them based on domain composition. This reclassification revealed not only a broader repertoire of potential RNA editing factors, but also extensive hallmark domain fusions among PPR, MORF, ORRM, and OZ proteins (except between MORF and OZ), mirroring their reported interactions^17,24,26,27,29^ and exemplifying the general link between domain fusion and protein-protein networking^76–78^. Based on these findings, we propose an evolutionary model where the composition of the editing machinery has progressively diversified from hornworts to angiosperms, likely contributing to the maintenance of efficient RNA editing (Extended Data Fig. 1). Collectively, these insights provide a refined view of the genomic innovations that have shaped the plant RNA machinery.

## Materials and Methods

### Collection and quality control of Archaeplastida genomes

To estimate the numbers of PPR, MORF, ORRM, and OZ genes and reconstruct their evolutionary landscapes, 364 Archaeplastida high-quality genomes were retrieved from public databases (Supplementary Table 1). To ensure accurate gene counts, only the longest protein isoform of each gene was retained. The completeness of annotated gene sets was estimated by Benchmarking Universal Single-Copy Orthologue (BUSCO) (version 5.8.2) with three datasets chlorophyta_odb10, viridiplantae_odb10, and eukaryota_odb10^79^. As these datasets contain abundant sequences of chlorophytes and angiosperms, genomes in these two lineages with BUSCO scores (“complete”) greater than 90% were retained. For other lineages, chromosome-level genomes with BUSCO scores (“complete”) greater than 85%; otherwise, the top 20% genomes with the highest BUSCO scores were retained within each lineage.

### Phylogenomic tree reconstruction of Archaeplastida

To reconstruct the phylogenomic tree of 364 Archaeplastida species, single-copy orthologous gene families were inferred by OrthoFinder (version 3.0.1b1)^80^, and 138 single-copy orthologous gene families covering at least 75% species were retained. These single-copy orthologous gene families were aligned by MAFFT (version 7.453)^81^ with the parameter “--auto”, and poorly aligned regions were identified and trimmed by BMGE (version 1.12) with BLOSUM30^82^. Gene trees were constructed by IQ-TREE (version 2.1.2)^83^, with the best-fit substitution model for each gene automatically selected according to Bayesian Information Criterion, and using parameters “-B 1000 -bnni -m MFP -T AUTO -ntmax 20”. The resulting trees were subsequently merged, and the species tree was inferred by ASTRAL (version 5.7.8)^84^ using *Cyanophora paradoxa* as the outgroup.

### Identification of PPR, MORF, ORRM, and OZ proteins

The hallmark domains and motifs of PPR (P, P1, P2, S1, S2, SS, L1, L2, E1, E2, E+, DYW, and DYW:KP), MORF (MORF), ORRM (RRM), and OZ (RanBP2 Znf) proteins were identified using hmmsearch in HMMER (version 3.2.1)^85^ with corresponding domain and motif profiles. PPR hallmark domain and motif profiles were obtained from a previous study^34^ and searched using hmmsearch with parameters “--domtblout -noali -E 0.1”. All hmmsearch scores for P, P1, P2, S1, S2, L1, L2, E1, E2, and E+ domains or motifs were greater than 0, with SS motif greater than 10, and DYW/DYW:KP domain greater than 30. As most PPR hallmark domain-containing proteins are localized to mitochondria and/or plastids^19,86^, they are defined as PPR proteins. In addition, PPR proteins were required to contain at least three of P, P1, P2, L1, L2, or SS motifs, unless they contained a DYW or DYW:KP domain.

The MORF domain profile, constructed using hmmbuild in HMMER (version 3.2.1) with nine MORF domains from *A*. *thaliana*, was searched using hmmsearch with parameters “--domtblout -noali -E 1e-10”. As MORF domain was homologous to PA domain, only hits with hmmsearch score greater than 36 were kept. The RRM domain profile was obtained from Pfam (PF00076.26)^87^ and searched using hmmsearch with parameters “--domtblout -noali -E 1e-10”. The hmmsearch score was greater than 0. GR domains, present in some MORF and ORRM proteins, were defined as regions containing at least eight glycines within a 20-amino-acid sliding window. The RanBP2 Znf profile was obtained from Pfam (PF00641.22) and searched using hmmsearch with the same parameters as for the RRM domain. As MORF, ORRM, and OZ proteins were localized to mitochondrion and/or plastids, their subcellular localization was predicted using DeepLoc (version 2.1)^88^ with “Accurate” mode, TargetP (version 2)^89^, Plant-mSub^90^ (https://bioinfo.usu.edu/Plant-mSubP), and LOCALIZER^91^ (https://localizer.csiro.au). Only localization results supported by at least two tools were retained. The domain composition was identified using InterPro (https://www.ebi.ac.uk/interpro)^92^.

### WGD Inference

To infer WGD, intra-genome alignments were conducted using BLASTP in the Basic Local Alignment Search Tool (BLAST, version 2.12.0+)^93^ with an E-value cutoff of 1e-5. Based on the alignments, homologous dotplots were generated using WGDI^94^, and synteny blocks were inferred using MCScanX^95^. Ks values of homologs in synteny blocks were calculated using KaKs_Calulator (version 3.0)^96^ and ParaAT (version 2.0)^97^ with parameters “-p proc -m mafft -f axt -g -k”. The median Ks value for each synteny block was calculated. Collectively, Ks peak counts and synteny blocks in the homologous dotplot were used to infer the number of WGD events.

### Identification and classification of duplicated gene pairs

To identify duplicated gene pairs, all protein sequences within a genome were aligned using BLASTP (BLAST, version 2.12.0+) with an E-value cutoff of 1e-5. The highest-scoring alignment for each gene was considered a duplicated pair. Alignments covering more than 60% of the longer gene (query or subject) were required to have a sequence similarity exceeding 20%. Alternatively, for alignments covering less than 60% of the longer gene, the alignment was required to span at least 105 amino acid residues with a sequence similarity exceeded 40%. Otherwise, the query and subject genes were considered singletons. Based on genomic loci, duplicated gene pairs were classified into four categories: tandem pairs (adjacent genes), proximal pairs (within 10 loci on the same chromosome), dispersed pairs (more than 10 loci apart on the same chromosome or on different chromosomes), and WGD pairs (located in synteny blocks).

### TE annotation

TEs were identified by a combination of *ab initio* and homology-based methods. For *ab initio* prediction, a consensus sequence library was built using RepeatModeler (version 2.0.6)^98^ with the parameter “-LTRStruct”. An LTR library was then constructed using LTR_Finder (version 1.07)^99^/LTR_FINDER_parallel (version 1.1)^100^, LTRharvest (version 1.6.6)^101^ with parameters “-minlenltr 100 -maxlenltr 7000”, and LTR_retriever (version 2.9.0)^102^. Both libraries were used to annotate TEs using RepeatMasker (version 4.1.8)^103^. For homology-based detection, TEs were identified using RepeatMasker (version 4.1.8) with Dfam (version 3.9)^104^ database. TE annotations obtained from both methods were then combined.

### Retroposition inference

Retroposition, underlying dispersed duplication gene pairs, was characterized by intron loss, polyA enrichment downstream of the 3’UTR, and flanking TEs. First, intron loss in one dispersed duplication gene pair was required to meet one of following criteria: (1) both genes had no intron; (2) a gene had no intron, while the other had at least one intron; (3) considering intron retention, a gene contained one intron, while the other had at least one intron; (4) both genes had more than one intron, differing by more than 10 introns. Second, polyA enrichment downstream of the 3’UTR of the intronless gene (or polyT enrichment upstream of the 5’UTR) was defined as at least eight adenines/thymines within a 10-base window. Genomic annotations were used to detect polyA/T enrichment in regions extending 1 kb downstream of the 3’UTR or upstream of the 5’UTR. If the 3’UTR or 5’UTR of a gene was not annotated, the shortest corresponding UTR in the genome was used. If no 3’UTR or 5’UTR was annotated in the genome, they were defined as the 100-bp regions downstream of the stop codon and upstream of the start codon, respectively. Third, dispersed duplication gene pairs with TEs located within 20 kb upstream and downstream were defined as TE-flanked.

### Identification of DYW and DYW:KP domain homologs and HGT inference

To explore HGT of plant DYW and DYW:KP domains, 10,407 protein sequences (hmmsearch score ≥ 30, length ≥ 130 aa) of 68 representative species were collected (Supplementary Table 8). DYW and DYW:KP domain homologs were searched in the NCBI Non-redundant Protein Sequence Database (NR) (acquisition date: 2025.01.19) using BLASTP (BLAST, version 2.12.0+) with an E-value cutoff of 1e-5. Protein sequences retrieved from NR originated from 40 lineages (Supplementary Table 9). Homologs exhibiting more than 20% sequence similarity to plant DYW and DYW:KP domains were further confirmed via hmmsearch in HMMER (version 3.2.1) using a score cutoff of 30. To avoid contamination, non-plant DYW and DYW:KP domains were retained only if present in at least two species within the corresponding lineage. Otherwise, non-plant DYW and DYW:KP domains detected in a single species were retained only if present in at least one additional genome assembly.

To reconstruct the phylogenetic tree of DYW and DYW:KP domains from plants and non-plants, domains from six representative plants (*P*. *pearsonii*, *T*. *lepidozioides*, *I*. *sinensis*, *A. spinulosa*, *G*. *montanum*, and *A*. *trichopoda*) and all non-plant counterparts were aligned using MAFFT (version 7.525), with poorly aligned regions trimmed by trimAl (version 1.5.rev0)^105^ using parameters “-gt 0.7 -cons 60 -st 0.1”. The phylogenetic tree was inferred using IQ-TREE (version 2.1.2) (Best-fit model: JTT + R10) with parameters “-B 1000 -bnni -m MFP -T AUTO --safe” and visualized with iTOL (version 7)^106^. Bootstraps were from 1,000 replicates. Multiple sequence alignment of DYW domains from plants and non-plants was performed using MEGA (version 11)^107^ and visualized with Jalview (version 2.11.4.0)^108^.

### Identification of MORF domain homologs

To identify homologs of plant MORF domains, 334 protein sequences (hmmsearch score ≥ 0, length ≥ 86 aa) from 33 representative species were collected (Supplementary Table 14). These MORF domain homologs searched in the NCBI NR using BLASTP (BLAST, version 2.12.0+) with an E-value cutoff of 1e-5. The protein sequences retrieved from NR originated from 45 lineages (Supplementary Table 9). Homologs with more than 20% similarity to plant MORF domains were retained. The MORF domains and their homologs were aligned using MAFFT (version 7.525), and the phylogenetic tree was constructed using FastTree^109^ (Model: Jones-Taylor-Thorton, CAT approximation with 20 rate categories) with parameters “-pseudo -gamma” and visualized by iTOL (version 7).

### Phylogenetic analysis of MORF domains

To reconstruct the phylogenetic tree of MORF domains, all MORF domains from gymnosperms and angiosperms were selected. These domains were aligned to MORF1∼MORF9 domains from *A*. *thaliana* to assign their types. Five PA domains from non-seed plants (Dicom.22G070000.1.p, HaB36G042, Phcar.S3G189600.t1, Phsp.C2G192800.t1, XP_024533311.1), exhibiting the highest similarity to MORF domains, were used as the outgroup. All MORF domains, along with the outgroup sequences, were aligned using MAFFT (version 7.525). The phylogenetic tree was inferred using IQ-TREE (version 2.1.2) (Best-fit model: JTT + R8) with parameters “-B 1000 -bnni -m MFP -T AUTO” and visualized by iTOL (version 7)^106^. Bootstraps were from 1,000 replicates.

### Analysis of PPR and MORF gene expression

To calculate gene expression, reference genome indexes were built using hisat2-build in HISAT2 (version 2.0.5)^110^ and rsem-prepare-reference in RSEM (version 1.3.1)^111^. Raw RNA sequencing data were quality-filtered using fastp (version 0.20.1)^112^ with parameters “-q 20 -u 40 -l 15 -g -x -r -W 4 -M 20 -w 12”. Strand specificity was inferred using infer_experiment.py in RSeQC (version 2.6.4)^113^. Reads mapping was conducted using STAR (version 2.7.1a)^114^. Gene expression matrix was generated using rsem-calculate-expression in RSEM (version 1.3.1) with parameters “--keep-intermediate-files --time --star -- append-names --output-genome-bam --sort-bam-by-coordinate”.

### Protein structure prediction and alignment

Protein structures were predicted using ColabFold (https://colab.research.google.com/github/sokrypton/ColabFold)^115^, and different protein structures were aligned using “Matchmaker” in ChimeraX (version 1.9)^116^.

### Data availability

All data used and generated in this study are available at the Open Archive for Miscellaneous Data (OMIX: OMIX012585)^117^ in the National Genomics Data Center (NGDC), China National Center for Bioinformation / Beijing Institute of Genomics, Chinese Academy of Sciences^118^.

### Code availability

The computational codes used in this study have been deposited in GitHub (https://github.com/Bioinfo-Ming/PlantEditosomeEvolution).

## ACKNOWLEDGEMENTS

We thank Shuhui Song, Lina Ma, Chao Zhang, Tianyi Xu, Tongtong Zhu, Dechang Yang, Shuangyang Wu, Lin Liu, Zhao Li, Yang Zhang, Xing Zheng, Yue Qi, Xinyu Zhou, Wenzhuo Cheng, Yuxin Qin, Miaomiao Wang, Shiting Wang, Lingjie Wang, Zihan Wang, Kehua Ma, Pan Li, Yiran Zhan, and Zheng Luo for their valuable discussions and advice. This work was supported by grants from National Natural Science Foundation of China [T2425005 and 32030021] and International Partnership Program of the Chinese Academy of Sciences [153F11KYSB20160008].

## CONTRIBUTOR INFORMATION

M.C. collected and analyzed the data, and wrote the manuscript. Z.Z. conceived, initiated and supervised this study. Q.M.Q. and Z.Z. revised the manuscript.

---

**Extended Data Fig. 1:**
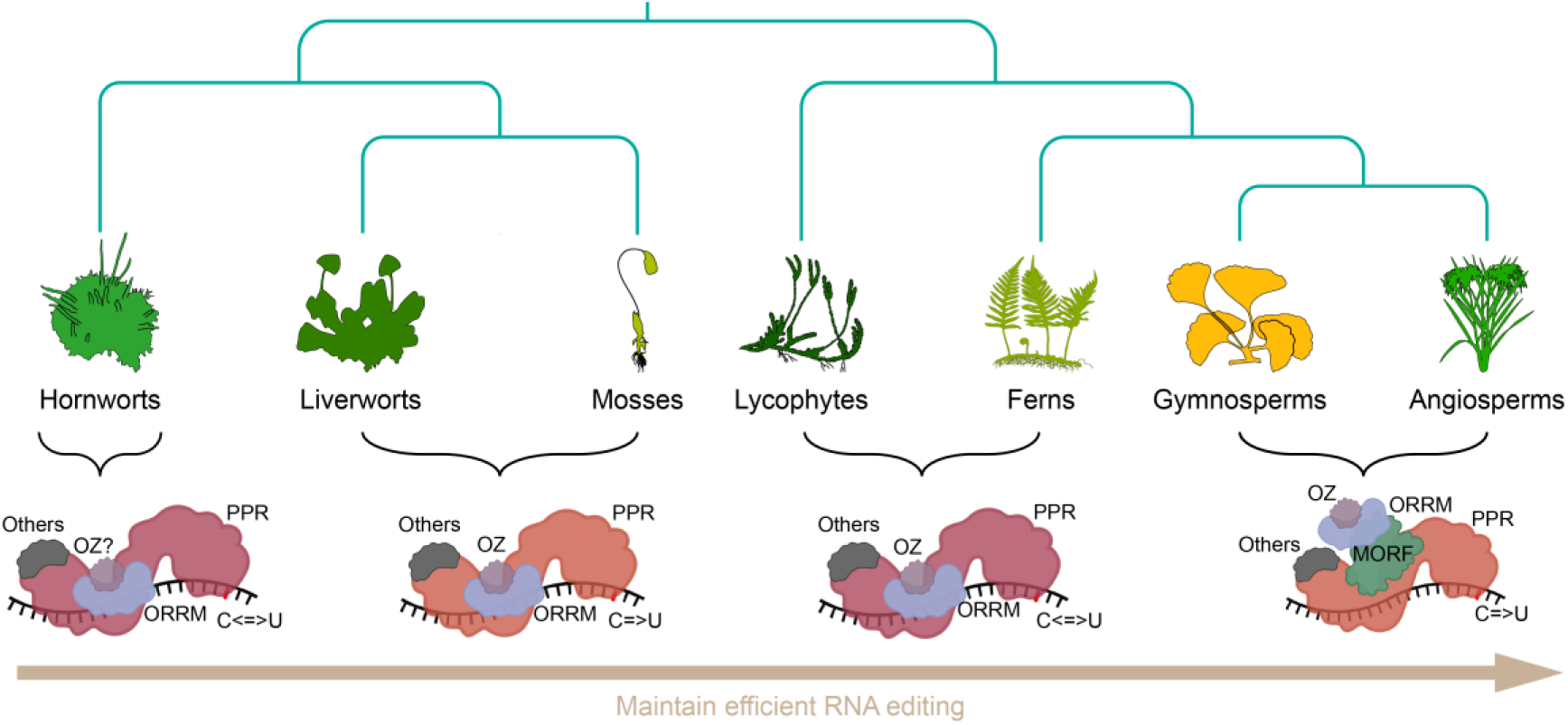
A proposed model for the evolutionary diversification of plant RNA editing factors. Editing factor composition varies from hornworts to angiosperms, potentially maintaining efficient RNA editing. The question mark indicates that OZ protein is rare in hornworts. Deep-rose PPR: DYW or DYW:KP subgroup PPR protein; red-orange PPR: DYW subgroup PPR protein. C=>U: C-to-U RNA editing; C<=>U: C-to-U or U-to-C RNA editing.

